# Health and disease imprinted in the time variability of the human microbiome

**DOI:** 10.1101/029991

**Authors:** Jose Manuel Martí, Daniel M. Martínez, César Gracia, Amparo Latorre, Andrés Moya, Carlos P. Garay

## Abstract

Human microbiota plays an important role in determining changes from health to disease. Increasing research activity is dedicated to understand its diversity and variability. We analyse 16S rRNA and whole genome sequencing (WGS) data from the gut microbiota of 97 individuals monitored in time. Temporal fluctuations in the microbiome reveal significant differences due to factors that affect the microbiota such as dietary changes, antibiotic intake, early gut development or disease. Here we show that a fluctuation scaling law describes the temporal variability of the system and that a noise-induced phase transition is central in the route to disease. The universal law distinguishes healthy from sick microbiota and quantitatively characterizes the path in the phase space, which opens up its potential clinical use and, more generally, other technological applications where microbiota plays an important role.

PACS numbers:

---

The desire to understand the factors that influence human health and cause diseases has always been one of the major driving forces of biological research. Modern high-throughput sequencing and bioinformatic tools provide a powerful means of understanding how the human microbiome contributes to health and its potential as a target for therapeutic interventions. High throughput methods for microbial 16S ribosomal RNA gene and WGS have now begun to reveal the composition of archaeal, bacterial, fungal and viral communities located both, in and on the human body. Biology has recently acquired new technological and conceptual tools to investigate, model and understand living organisms at the system level, thanks to the spectacular progress in quantitative techniques, large-scale measurement methods and the integration of experimental and computational approaches. Systems Biology has mostly been devoted to the study of well-characterized model organisms but, since the early days of the Human Genome Project it has become clear that applications of system-wide approaches to Human Biology would bring huge opportunities in Medicine. Here we present the imprints of disease in macroscopic properties of the system, by studying the temporal variability in the microbiome.

To this end, we have analysed more than 35000 time series of taxa from the gut microbiomes of 97 individuals (sampling from three up to 332 time points), obtained from publicly available high throughput sequencing data on: healthy individuals over a long term span[1], people with various degrees of obesity[2], twin pairs discordant for kwashiorkor[3], response to diet changes[4] or to antibiotic perturbation[5], and subjects diagnosed with irritable bowel syndrome (IBS)[6]. We engineered a complete software framework and a web platform, ComplexCruncher, ready to be implemented by other users.

We summarize our dataset in ST1-6 (Tables 1-6 in suplemental material). The bacteria and archaea taxonomic assignations were obtained by analysing 16S rRNA sequences, which were clustered into operational taxonomic units (OTUs) sharing 97 % sequence identity using QIIME[7]. WGS data[3] were analysed and assigned at strain level by the Livermore Metagenomic Analysis Toolkit (LMAT)[8], according to their default quality threshold. Genus, with best balance between error assignment and number of taxa, was chosen as our reference taxonomic level. We have verified that our conclusions are not significantly affected by selecting family or species as the reference taxonomic level (see Figure 1 in supplemental material).

**FIG. 1:**
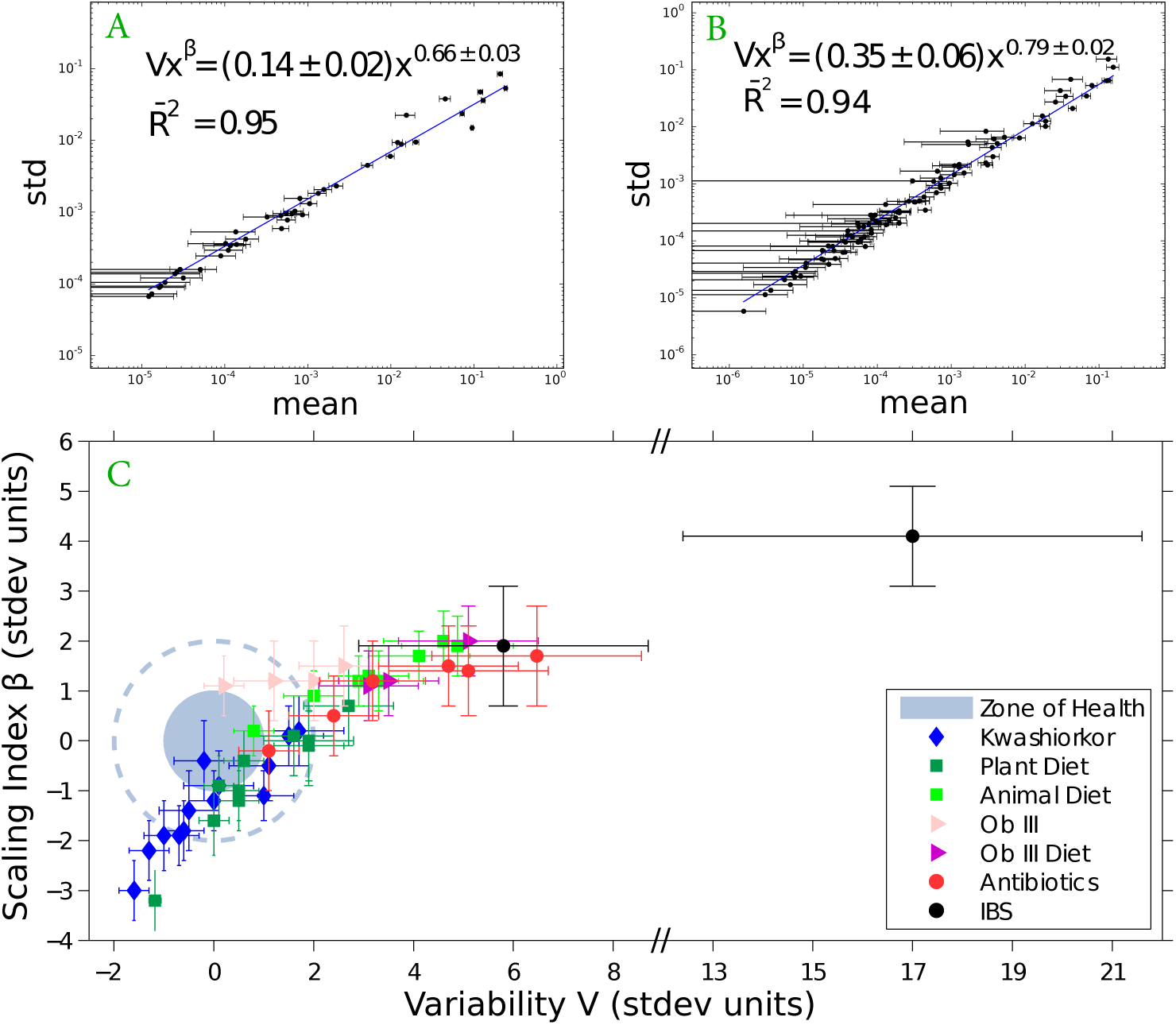
Taylor’s law parameter space. X-weighted power-law fits of the standard deviations versus the mean values for each bacterial genus monitored in time. We show the fit for samples from a healthy subject (Figure A) and from a subject diagnosed with irritable bowel syndrome (Figure B) studied in our lab[6]. We have compiled all data studied in this work in Figure C. The coloured region (dashed line) corresponds to 68% (95%) CL region of healthy individuals in the Taylor parameter space. Points with errors place each individual gut microbiome in the Taylor space. Note that the parameters have been standardized (stdev units) to the healthy group in each study for demonstrative purposes.

We have analysed the microbiome temporal variability to extract global properties of the system. As fluctuations in total counts are plagued by systematic errors we worked on temporal variability of relative abundances for each taxon. Our first finding was that, in all cases, changes in relative abundances of taxa follow a universal pattern known as the fluctuation scaling law[9] or Taylor’s power law[10], i.e., microbiota of all detected taxa follows a power law dependence between mean relative abundance *x_i_* and dispersion 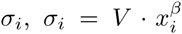. While the law is universal, spanning six orders of magnitude in the observed relative abundances, the power law (or scaling) index *β* and the variability *V* (hereafter Taylor parameters) appear to be correlated with the stability of the community and the health status of the host, which is the main finding in this letter (see Figure 1). Taylor parameters describing the temporal variability of the gut microbiome in our sampled individuals are shown in ST1-6. Our results hint at a universal behaviour. Firstly, the variability (which corresponds to the maximum amplitude of fluctuations) is large, which suggests resilient capacity of the microbiota, and the scaling index is always smaller than one, which means that, more abundant taxa are less volatile than less abundant ones. Secondly, Taylor parameters for the microbiome of healthy individuals in different studies are compatible within estimated errors. This enables us to define the health zone in the Taylor parameter space. We can better visualize the results of individuals from different studies by standardizing their Taylor parameters, where standardization means that each parameter is subtracted by the mean value and divided by the standard deviation of the group of healthy individuals in the study (see Section 12 and ST1-6). The zone of health and the standardized Taylor parameters for individuals whose gut microbiota is threatened (i.e., suffering from kwashiorkor, altered diet, antibiotics, IBS) is shown in Figure 1. Children developing kwashiorkor show smaller variability than their healthy twins. A meat/fish-based diet increases the variability significantly when compared to a plant-based diet. All other cases presented increased variability, which is particularly severe, and statistically significant at more than 95% CL, for obese patients grade III on a diet, individuals taking antibiotics or IBS-diagnosed patients. A global property emerges from all worldwide data collected: Taylor parameters characterize the statistical behaviour of microbiome changes. We have verified that our conclusions are robust to systematic errors due to taxonomic assignment.

Taylor’s power law has been explained in terms of various effects, all without general consensus. It can be shown to have its origin in a mathematical convergence similar to the central limit theorem, so virtually any statistical model designed to produce a Taylor law converge to a Tweedie distributional], providing a mechanistic explanation based on the statistical theory of errors[12–14]. To unveil the generic mechanisms that drive different scenarios in the *β*–V space, we model the system by assuming that taxon relative abundance follows a Langevin equation with a deterministic term that captures the fitness of each taxon and a randomness term with Gaussian random noise[15]. Both terms are modelled by power laws, with coefficients that can be interpreted as the taxon fitness *F_i_* and the variability *V* (see Section 1 in supplemental material). When *V* is sufficiently low, abundances are stable in time. Differences in variability *V* can induce a noise-induced phase transition in relative abundances of taxa. The temporal evolution of the probability of a taxon having abundance *x_i_* given its fitness is governed by the Fokker–Planck equation. The results of solving this equation show that the stability is best captured by fitness F and amplitude of fluctuations V phase space (see Figure 2). The model predicts two phases for the gut microbiome: a stable phase with large variability that permits some changes in the relative abundances of taxa and an unstable phase with larger variability, above the phase transition, where the order of abundant taxa varies significantly with time. The microbiome of all healthy individuals was found to be in the stable phase, while the microbiome of several other individuals was shown to be in the unstable phase. In particular, individuals taking antibiotics and IBS–diagnosed patient P2 had the most severe symptoms. In this phase diagram, each microbiota state is represented by a point at its measured variability V and inferred fitness F. The model predicts high average fitness for all taxa, i.e., taxa are narrowly distributed in F. The fitness parameter has been chosen with different values for demonstrative purposes. Fitness is larger for the healthiest subjects and smaller for the IBS–diagnosed patients.

**FIG. 2:**
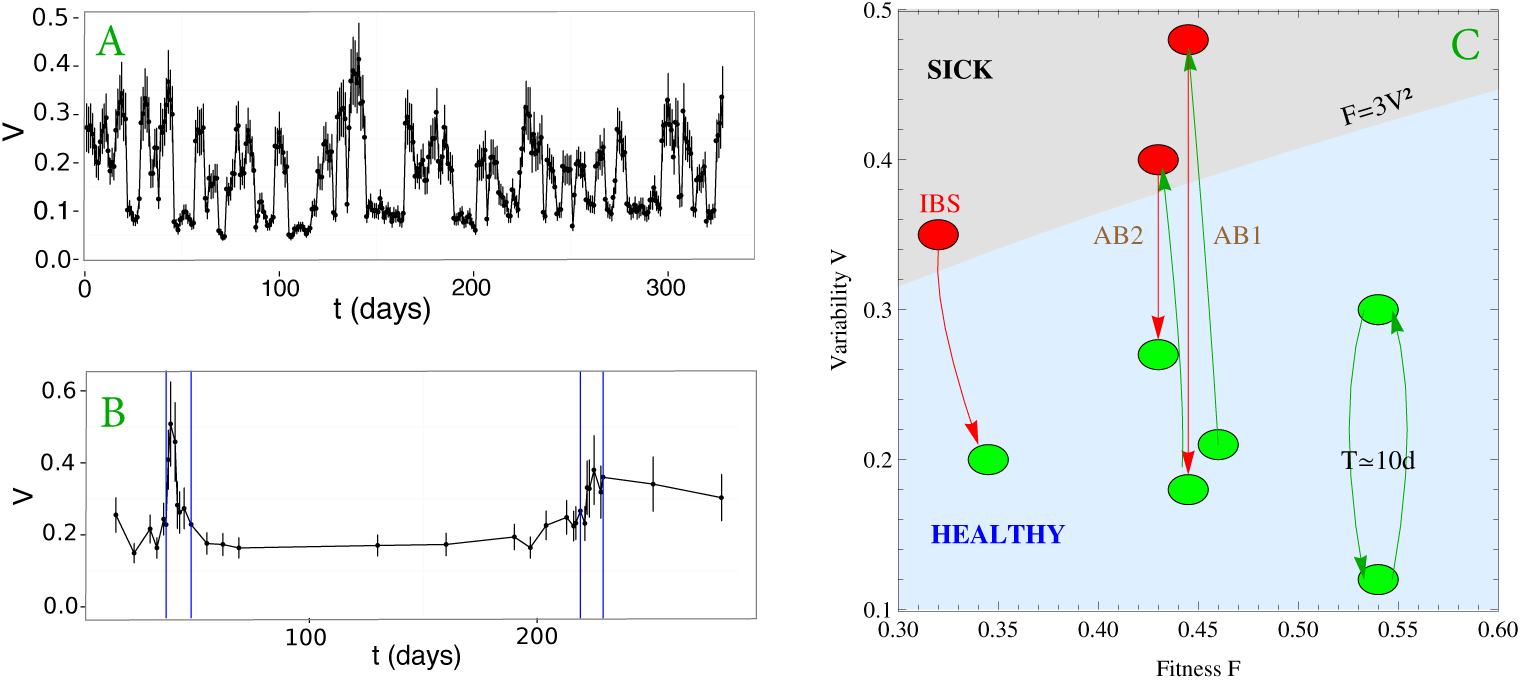
Phase space of microbiota stability. We show the Variability parameter as a function of time for the male in Caporaso’s study[1] (Figure A) and for the patient D in the antibiotics study[5] (Figure B). Periods of antibiotic treatment are shown by blue vertical lines. Microbiota states can be placed in the phase space F-V (Figure C). The light blue shaded region corresponds to the stable phase, while the grey shaded region is the unstable phase (the phase transition line is calculated for *α* = *β* = 0.75). We place healthy individuals (green) and individuals whose gut microbiota is threatened (antibiotics, IBS) in the phase space fitness - variability. Gut microbiota of healthy individuals over a long term span show a quasi-periodical variability (central period is ten days). We show that taking antibiotics (AB1 and AB2 correspond to first and second treatment respectively) induces a phase transition in the gut microbiota, which impacts its future changes. We also show an IBS–diagnosed patient transiting from the unstable to the stable phase.

In summary, we have quantitatively characterized whether the microbiota belongs to a healthy individual or a subject corresponding to an altered or pathological state (ie, altered diet, antibiotic treatment, early gut development, diagnosed IBS). Deciphering the mechanisms of disease requires in depth knowledge of the underlying biological mechanisms. We describe here the macroscopic behavior of disease by a noise-induced phase transition with a control parameter that can be measured by the temporal variability of the microbiome. The microbiota of healthy individuals and of individuals with pathologies represent different phases separated by this noise-induced phase transition. Improved high–throughput sequencing of samples from individuals monitored over time and taxonomic assigning methods will provide a better distinction among pathologies or altered states of the microbiota.

## Acknowledgements.

— This work is supported by Generalitat Valencia Prometeo Grants II/2014/050, II/2014/065, by the Spanish Grants FPA2011-29678, BFU2012-39816-C02-01 of MINECO and by PITN-GA-2011-289442-INVISIBLES. JMM and DMM acknowledge FPI and FISABIO fellowships.

